# Infectious bronchitis virus vaccination, but not the presence of XCR1, is correlated with large differences in chicken caecal microbiota

**DOI:** 10.1101/2024.05.22.595109

**Authors:** Laura Glendinning, Zhiguang Wu, Lonneke Vervelde, Mick Watson, Adam Balic

## Abstract

The chicken immune system and microbiota play vital roles in maintaining gut homeostasis and protecting against pathogens. In mammals, XCR1+ conventional dendritic cells (cDCs) are located in the gut-draining lymph nodes and play a major role in gut homeostasis. These cDCs sample antigens in the gut luminal contents and limit the inflammatory response to gut commensal microbes by generating appropriate regulatory and effector T-cell responses. We hypothesised that these cells play similar roles in sustaining gut homeostasis in chickens, and that chickens lacking XCR1 were likely to contain a dysbiotic caecal microbiota. Here we compare the caecal microbiota of chickens that were either heterozygous or homozygous XCR1 knockouts, that had or had not been vaccinated for infectious bronchitis virus (IBV). We used short-read (Illumina) and long-read (PacBio HiFi) metagenomic sequencing to reconstruct 670 high-quality, strain-level metagenome assembled genomes. We found no significant differences between alpha-diversity or the abundance of specific microbial taxa between genotypes. However, IBV vaccination was found to correlate with significant differences in the richness and beta-diversity of the microbiota, and to the abundance of forty bacterial genera. In conclusion, we found that a lack of XCR1 was not correlated with significant changes in the chicken microbiota, but IBV vaccination was.

**Data Summary:** The raw sequencing data for this project, as well as primary assemblies, putative genome bins and species-level MAGs, are available in the European Nucleotide Archive under project PRJEB64517. Strain-level MAGs are available through Edinburgh DataShare (https://doi.org/10.7488/ds/7678).

**Impact statement:** Chickens play a vital role in global food systems, with 74 billion chickens killed for meat and 1.6 trillion chicken eggs produced in 2021 alone. The gut microbiota plays a vital role in the health and nutrition of the chicken, contributing to gut homeostasis and the production of nutrients that can be absorbed and used by the host bird. The chicken gut microbiota represents an excellent target to improve pathogen resistance and nutrition, as farmed chickens are not exposed to a maternal hen and thereby develop a low-diversity gut microbiota that negatively impacts their immune development and gut health. In order to develop such microbiota interventions, we first need to understand the fundamental biology behind immune-microbiota interactions. While it is well known that the chicken gut microbiota plays an important role in gut homeostasis, the mechanisms behind this phenomenon are poorly understood. In mammals, XCR1+ conventional dendritic cells have been shown to play a role in maintaining gut homeostasis. In this study we compare the gut microbiota of heterozygous and homozygous XCR1 knockout chickens that did or did not receive infectious bronchitis virus vaccination. We found that the gut microbiota of heterozygous and homozygous XCR1 knockout chickens did not differ significantly, but IBV vaccination did significantly correlate with differences in the microbiota composition.

## Introduction

The chicken immune system and gut microbiota both play vital roles in maintaining gut homeostasis and protecting against pathogens. It is well known that the immune system plays a crucial role in shaping and maintaining the chicken gut microbiota, and that the gut microbiota itself contributes to immune system maturation (1, 2). Raising chickens in a germ-free environment has been shown to lead to decreased maturation of both the innate and adaptive immune response, and gut physiology (2, 3). Providing chicks with exposure to caecal/faecal content transplants from adult birds, or with synthetic communities of microbes constructed from microbes isolated from the chicken gut, has been shown to modulate the immune system (4–6). Similar microbiota transplants have also been shown to have a protective effect against gut pathogens such as *Salmonella* spp. (5, 7, 8).

Mammalian XCR1+ conventional dendritic cells (cDCs) are located in the gut-draining lymph nodes and play a major role in gut homeostasis (9). In chickens, which lack draining lymph nodes, XCR1+ cDCs are found throughout the intestinal tissues (10). cDCs sample antigens in the gut luminal contents and limit the inflammatory response to gut commensal microbes by generating appropriate regulatory and effector T-cell responses (11). XCR1 is a chemokine receptor exclusively expressed by cDCs in chickens and a subset of cDCs in mammals (10, 12). Mice lacking XCR1 show decreased intraepithelial and lamina propria T-cells and an increased susceptibility to gut inflammation (13). Mice lacking XCR1+ cDCs also display an altered immune response to a common gut commensal (*Cryptosporidium tyzzeri*), correlated with opportunistic infection by the microbe (14). We hypothesised that these cells would play similar roles in sustaining gut homeostasis in chickens, and therefore chickens lacking XCR1 were likely to contain a dysbiotic caecal microbiota. Increased gut inflammation in particular may lead to a decrease in bacteria belonging to the phylum Firmicutes (15), which has previously identified as one of the most common phyla in chickens (16, 17). Increased gut inflammation also commonly leads to an increase in opportunistically pathogenic bacteria (18).

In this study we compare the microbiota of wild-type and XCR1 knockout chickens, that have or have not been vaccinated for infectious bronchitis virus (IBV). It has previously been shown using 16S rRNA gene sequencing that the caecal microbiota of chickens differs according to both IBV vaccination and genetic line, after infection with IBV (19). We use shotgun metagenomic sequencing to characterise the gut microbiota, and to test whether genetic line and IBV vaccination without subsequent IBV infection correlate with differences in the gut microbiota.

## Methods

### Sample collection

All birds were hatched and housed in premises licensed under a UK Home Office Establishment License in full compliance with the Animals (Scientific Procedures) Act 1986 and the Code of Practice for Housing and Care of Animals Bred, Supplied or Used for Scientific Purposes. Production of the XCR1 knockout line (10) was carried out under UK Home Office Licenses (70/8528; 70/8940 and PP9565661). Ethical approvals were obtained from The Roslin Institute’s and University of Edinburgh’s Animal Welfare Ethics Review Board and conducted under the authorisation of a UK Home Office Project License (PE263A4FA) and adhered to the guidelines and regulations of the UK Home Office ‘Animals (Scientific Procedures) Act of 1986’.

All birds were obtained from the National Avian Research Facility at The Roslin Institute, University of Edinburgh. Twenty-four chickens, heterozygous for the XCR1-iCaspase9-RFP transgene (n=12) or homozygous for the XCR1-iCaspase9-RFP transgene (n=12) (10), were either vaccinated intranasally and intraocularly with 3 doses live attenuated IBV vaccine (Nobilis IB Ma5) at 8 weeks old and boosted at 18 weeks old, or were controls receiving PBS (**Table 1**). Previously we have shown that birds homozygous for the *iCaspase9-RFP* transgene are deficient (“Knock-out”: “KO”) in XCR1 expression, whereas heterozygous birds exhibited the same level of expression as non-transgenic wild-type (WT) birds (10). Heterozygous birds are therefore functionally wild-type for XCR1 expression and are suitable for assessing XCR1^+^ cDC development and function. Hereafter, groups of heterozygous and homozygous birds are referred to as “XCR1 WT” and “XCR1 KO” respectively.

**Table 1:** Metadata for chickens used in study.

All chickens were fed a standard commercial broiler diet. All birds were housed in the same room until 8 weeks of age, after which vaccinated and control birds were housed in separate rooms to prevent spread of the live vaccine to control birds. Vaccinated birds were culled 7 days after the IBV booster and control birds were culled 8 days after the IBV booster. Caecal contents were collected, snap frozen and stored at -20°C.

### DNA extraction and sequencing

DNA was extracted from caecal contents using the QIAamp PowerFecal Pro DNA Kit (Qiagen) according to the manufacturer’s instructions, with bead beating in a FastPrep homogeniser for 5 metres/second for 40 seconds. RNA was then removed by incubating 50 µl of each sample with 5 µl of RNase Cocktail Enzyme Mix (Thermo Fisher Scientific) for 1 hour at room temperature. DNA was purified using a 1:1 ratio of sample to AMPure XP Beads (Beckman Coulter), and eluted into EB buffer. Short read sequencing was conducted on all samples, by Novogene Sequencing (Novogene Corporation Inc), using a NovaSeq producing paired-end 150 bp reads. Long-read sequencing was performed on one sample per group (n = 4) by Edinburgh Genomics, using a PacBio Sequel IIe system producing HiFi reads.

### Quality control of reads

Adaptors were trimmed from Illumina reads using Trimmomatic (20) (v. 0.36, options: LEADING:3 TRAILING:3 MAXINFO:40:0.5 AVGQUAL:30 MINLEN:36). The host genome (*Gallus gallus*: GCF_000002315.6) and feed genomes (*Glycine max*: GCF_000004515.6, *Aegilops tauschii subsp. Strangulate:* GCF_002575655.2, *Triticum aestivum*: GCF_018294505.1, *Zea mays*: GCF_902167145.1, *Hordeum vulgare subsp. Vulgare*: GCF_904849725.1) were downloaded from RefSeq, then feed and host reads were removed from adaptor trimmed read files. Illumina reads were mapped to host and feed reference genomes using BWA-MEM (21) (v. 0.7.17), followed by SAMtools (22) (v. 1.17, samtools fastq -f 12) to select reads where both paired-end reads were unmapped. HiFi reads were mapped to host and feed reference genomes using Minimap2 (23) (v.2.24, options: - ax map-hifi) followed by SAMtools (samtools fastq -f 4) to select reads that were unmapped. Quality controlled reads were used for all further taxonomic analysis and genome assembly steps.

### Taxonomic assignment of reads

Kraken2 (24) (v. 2.1.2) was used to assign taxonomy to host and feed filtered reads. First, “kraken2-build --download-taxonomy” was used to download the NCBI taxonomy (download date: 24^th^ April 2023). Feed/host genomes and GenBank microbial genomes downloaded from previous chicken microbiota studies (NCBI BioProjects: PRJNA715658, PRJEB33338, PRJNA543206, PRJNA377666 and PRJNA668258: downloaded 2^nd^ May 2023) were added to the Kraken2 database using “kraken2-build --add-to-library”. Standard Kraken2 genomes were downloaded using “kraken2-build --download-library” (bacteria, archaea, plasmid, viral, fungi, plant, protozoa: downloaded April 24^th^ 2023). The database was then built using “kraken2-build –build”. Reads were classified by Kraken2 against this database, using the option “--paired“ for Illumina reads.

### Primary assembly and genome bin construction

For single-sample assembly and coassembly, Illumina reads were assembled into contigs using Megahit (25) (v.1.2.9, options: --kmin-1pass --k-list 27,37,47,57,67,77,87 --min-contig-len 1000). Illumina reads from all samples were mapped separately to each assembly using BWA-MEM. From the resulting BAM files, depth files were created using the command jgi_summarize_bam_contig_depths from metabat2 (26) (v.2.15). Using these depth files, Metabat2 was used to construct genome bins.

For single-sample assembly and assembly of pooled HiFi reads, we used three different assemblers to maximise the number of bins obtained from HiFi data: Hifiasm Meta (27) (v.hamtv0.3.1, options: -oasm), metaFlye (v2.9.2, flye --meta --pacbio-hifi) and HiCanu (28) (v.2.2, options: maxInputCoverage=100000 correctedErrorRate=0.105 genomeSize=5m batMemory=200 (increased to 300 for sample 6737_HOM_CTRL) useGrid=false -pacbio-hifi). HiFi reads from all samples were mapped separately to each assembly using Minimap2 (-ax map-hifi). From the resulting BAM files, depth files were created using the command jgi_summarize_bam_contig_depths from metabat2. Using these depth files, Metabat2 was used to construct putative genome bins. For each assembly, circular contigs and contigs >0.5mb were also output into separate fastas and classed as putative genome bins.

For hybrid assembly of single samples, metaSPAdes (29) (SPAdes v.3.15.5) was used. BWA-MEM was used to map Illumina reads from a single sample back to its own assembly. From the resulting BAM file, depth files were created using the command jgi_summarize_bam_contig_depths from metabat2. Using these depth files, Metabat2 was used to construct genome bins. As for assemblies constructed only from HiFi reads, contigs >0.5mb were also output into separate fastas and classed as putative genome bins. No circular contigs were identified in the hybrid assemblies (visualised using Bandage (30) (v.0.8.1)).

### Quality control, dereplication and quantification of genome bins

For each HiFi and hybrid assembly, bins originating from that assembly were processed using DAS Tool (31) (v.1.1.6) to create a set of non-redundant bins. Firstly, the script Fasta_to_Contigs2Bin.sh from the DAS Tool suite was used to convert genome bins into a contigs-to-bin table. Then DAS Tool was ran using the options --write_bin_evals --write_bins. Checkm2 predict (32) (v.1.0.2) was used to check the quality of bins output by DAS Tool (HiFi and hybrid assemblies) and metabat2 (Illumina assemblies). These bins were used as input for the steps below.

Using dRep (v.3.4.3, options: -comp 80 -con 10), bins were dereplicated at 99% (--S_ani 0.99) average nucleotide identify (ANI) to construct strain-level metagenome assembled genomes (MAGs), and at 95% (--S_ani 0.95) ANI to construct species-level MAGs. Average amino acid identity (AAI) between MAGs was calculated using CompareM (33) (v.0.1.2). CoverM genome (34) (v.0.6.1) was used to quantify the abundance of MAGs in samples (methods: mean, relative_abundance, trimmed_mean, covered_bases, variance, length, reads_per_base, count, rpkm and tpm), with BWA-MEM for Illumina sequenced samples (options: -p bwa-mem --min-read-aligned-percent 75 --min-read-percent-identity 95 --min-covered-fraction) and Minimap2 for HiFi samples (options: -p minimap2-no-preset -- minimap2-parameters ’-ax map-hifi’ --min-read-aligned-percent 75 --min-read-percent-identity 95 --min-covered-fraction 0).

### MAG taxonomic and functional annotation

GTDB-Tk (35) (v2.3.0) and its dependencies (36–41) were used to assign taxonomy to MAGs. The Genome Database Taxonomy (42) (GTDB) was downloaded on 5^th^ July 2023, and used as a reference database for the command gtdbtk classify_wf. Phylophlan (43) (v.3.0.3, options: -d phylophlan --diversity high --subsample Phylophlan) was used to construct taxonomic trees, using the supermatrix_aa.cfg database (downloaded 10^th^ July 2023). Trees were rerouted at the branch between Archaea and Bacteria using FigTree (44) (v.1.4.3). Itol.toolkit (45) (v.1.1.5) was used to produce Interactive Tree Of Life format annotation files. Interactive Tree Of Life (46) (v.6) was used to visualise trees. AMR genes were identified using RGI (47) (v.6.0.2) and the Comprehensive Antibiotic Resistance Database (downloaded 10^th^ July 2023). METABOLIC (48) and its associated databases (GTDB (42), KOfam (49), Pfam (50), MEROPS (51), TIGRFAMs (52), dbcan2 (53) and a custom METABOLIC HMM) were downloaded on the 12^th^ February 2024.

### Statistics and graphical analyses

R packages used in our analysis included: ANCOMBC (54) (v.2.0.3), cowplot (55) (v. 1.1.1), dplyr (56) (v.1.1.1), ggplot2 (57) (v. 3.4.2), phyloseq (58) (v.1.42.0), reshape2 (59) (v.1.4.4), tidyr (60) (v.1.3.0), tidyverse (61) (v. 2.0.0) and vegan (62) (v.2.6-4). The relative abundance of MAGs in samples, was calculated as Transcripts Per Million (TPM), as output by CoverM. TPM is commonly used in RNA-seq studies, but is also useful for calculating MAG abundance (63–65) as it corrects for library size and genome length. TPM values are reported in the text as mean ± standard deviation. Using the vegan package, Bray-Curtis dissimilarity values were used to construct NMDS graphs, and to conduct PERMANOVA analysis (adonis) with sex, genotype and IBV vaccination included as factors, with their interactions. Alpha diversity values were calculated using the vegan package: the inverse Simpson’s value was used for diversity and the Chao1 index for richness. Differences in alpha diversity between groups were assessed using the Kruskal-Wallis rank sum test. Further pairwise tests were conducted using the Wilcoxon rank sum exact test with Bonferroni corrections for multiple tests.

Taxa that were significantly differently abundant between groups were identified using ANCOMB-BC2 (66) from the ANCOMBC package. TPM is not appropriate as input for ANCOM-BC2, as it requires “raw” count values. However, by only providing raw counts we do not control for genome size. Genome size corrected “raw” count values were produced by transforming TPM values into relative abundances and then multiplying these values by the total raw read counts (as output by CoverM) for each sample and rounding to the nearest integer. These count values were then used as input for ANCOM-BC2 (options: p_adj_method = “hochberg”, prv_cut = 0.10).

### Data availability

The raw sequencing data for this project, as well as primary assemblies, putative genome bins and species-level MAGs, are available in the European Nucleotide Archive under project PRJEB64517. Strain-level MAGs are available through Edinburgh DataShare (https://doi.org/10.7488/ds/7678).

## Results

### Sequence quality

We collected caecal samples from chickens that were either heterozygous or homozygous for the iCaspase9-RFP transgene, half of which had been vaccinated for IBV. Samples originated from four groups: XCR1 WT vaccinated with IBV (n=6), XCR1 KO vaccinated with IBV (n=6), XCR1 WT not vaccinated with IBV (n=6), and XCR1 KO not vaccinated with IBV (n=6). Extracted DNA from all samples underwent Illumina sequencing (150 bp reads), and DNA from one sample per group also underwent PacBio HiFi sequencing. After adaptor trimming but prior to contamination removal, HiFi samples contained 1,679,798 ± 316,552 reads (N50: 9365 ± 276 bp) and Illumina samples contained 45,567,257 ± 5,532,248 reads. After removal of host DNA and DNA expected to originate from feed sources, HiFi samples contained 1,581,201 ± 312,674 reads (N50: 9484 ± 283 bp) and Illumina samples contained 39,104,307 ± 5,885,045 reads. Reads were assembled into contigs then binned into putative genome bins. Dereplication and quality filtering of bins resulted in 480 (species-level: ANI 95%) and 670 (strain-level: ANI 99%) metagenome assembled genomes (MAGs) (**Figure 1**, **Table 2**). Of the 670 strain-level MAGs, 123 originated from single-sample HiFi assemblies, 112 from the pooled HiFi assembly, 244 from single-sample Illumina assemblies, 162 from the Illumina coassembly, and 29 from the hybrid assemblies. At species-level, 74 single-contig MAGs were produced from long-read or hybrid assembly, including 28 closed genomes.

**Figure 1:**
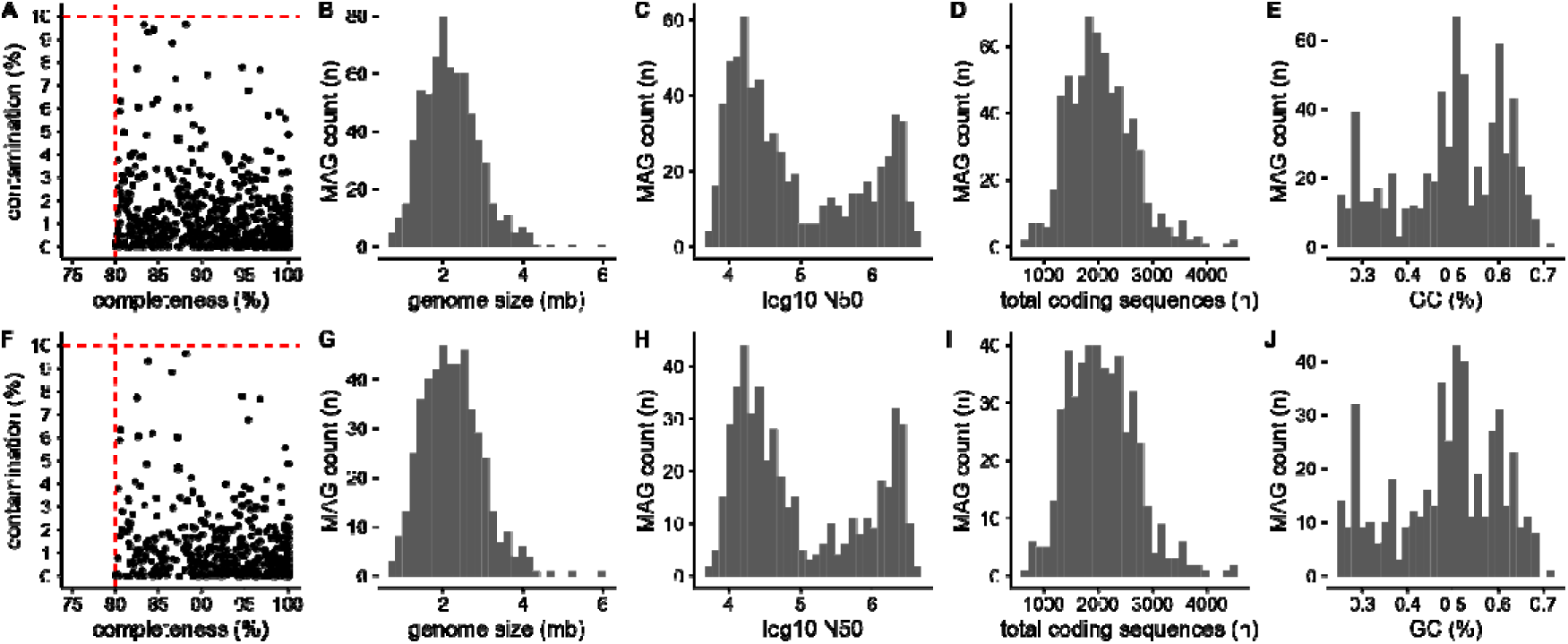
Genome statistics of non-redundant strain-level (A-E) and species-level (F-J) metagenome assembled genomes. A and F: Completeness and contamination – dashed red lines indicate cut-offs for defining genome bins as metagenome assembled genomes. B and G: Total size of each genome (mb). C and H: log10 N50 per genome. D and I: Total coding sequences per genome (n). E and J: Percentage GC content of each genome.

**Table 2:** Description of each chicken species-level MAG (metagenome-assembled genome), including GTDB-Tk taxonomy, and CheckM2 completeness, contamination and genome quality information.

### Taxonomy of MAGs and reads

Species-level MAGs originated from a wide diversity of taxa, but were dominated by members of the Bacillota phyla (**Figure 2**, **Table 2**). The vast majority of MAGs originated from bacteria, with only one MAG (*Methanobrevibacter_A woesei*) originating from an Archaeon. The most abundant phylum amongst our MAGs was Bacillota_A (n = 271), which predominantly contained members of the orders Oscillospirales (n = 88), Lachnospirales (n = 82) and Christensenellales (n = 52). This was also on average the most abundant phylum in samples (**Figure 3**) (388,536 ± 69,442 TPM). The second most abundant phylum amongst our MAGs was Bacillota (n =57), which mainly contained members of the RF39 (n = 27) and Lactobacillales (n = 17) orders. The phylum Bacteroidota (n= 56) was almost as abundant amongst our MAGs as Bacillota, and contained little diversity at order level, with only one MAG not originating from Bacteroidales. Despite containing similar numbers of MAGs, the average abundance of the Bacteroidota in our samples (279,451 ± 56,295 TPM) was much greater than that of the Bacillota (81,277 ± 42,236 TPM). Also, despite containing less MAGs than the above phyla (n = 34), the Actinomycetota were the third most abundant phyla in our samples (102,033 ± 44,402). The metabolic potential of MAGs varied greatly (**Table 3**), and nearly all MAGs contained at least one AMR gene (n=458: **Table 4**) with the vast majority of AMR genes targeting glycopeptide antibiotic (75% AMR genes).

**Figure 2:**
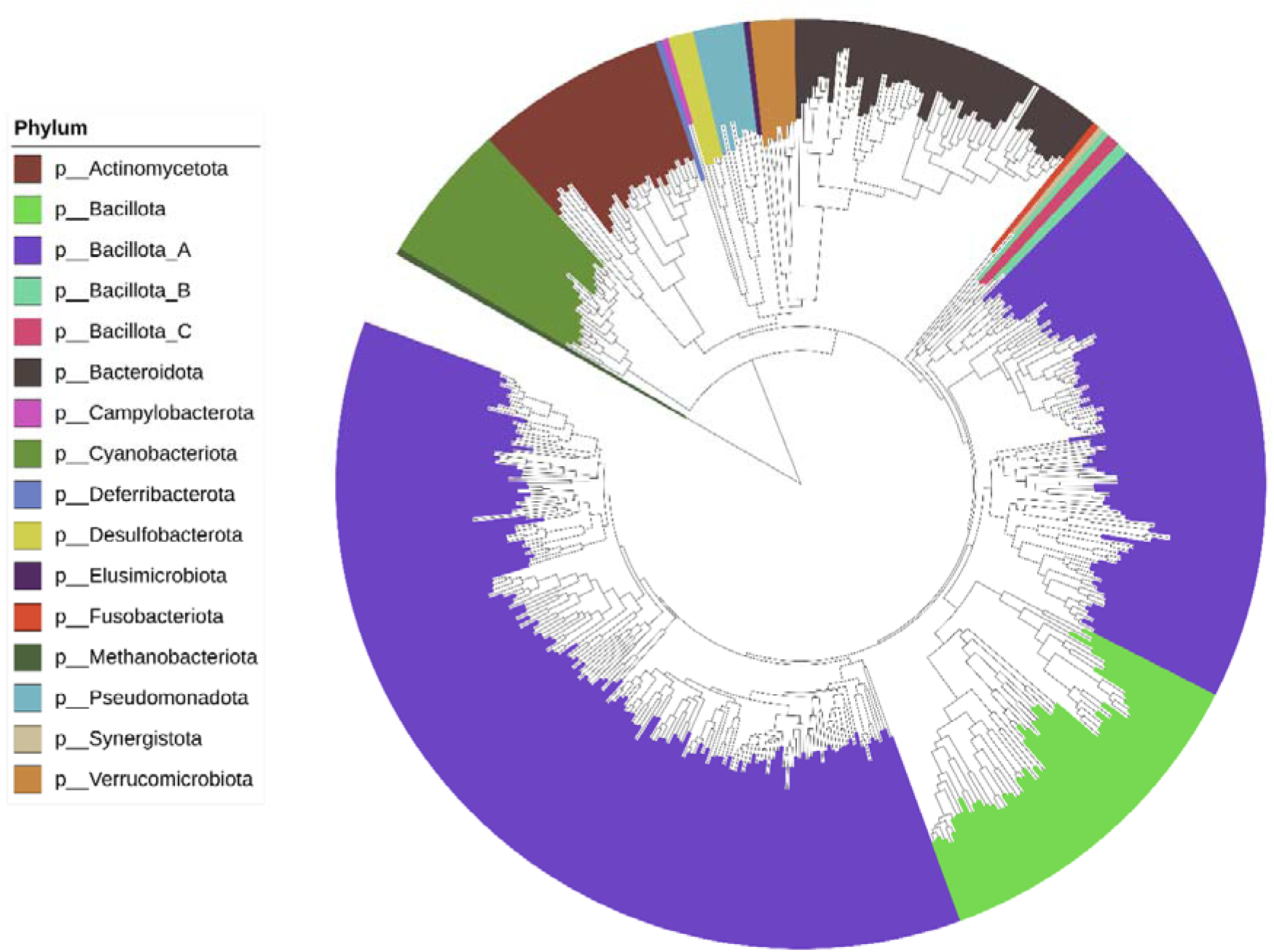
Taxonomic tree displaying the phylogeny of species-level MAGs, with tree rooted between Archaea and Bacteria. Only the 441 MAGs that included at least 100 universal proteins, as defined by Phylophlan, are included.

**Figure 3:**
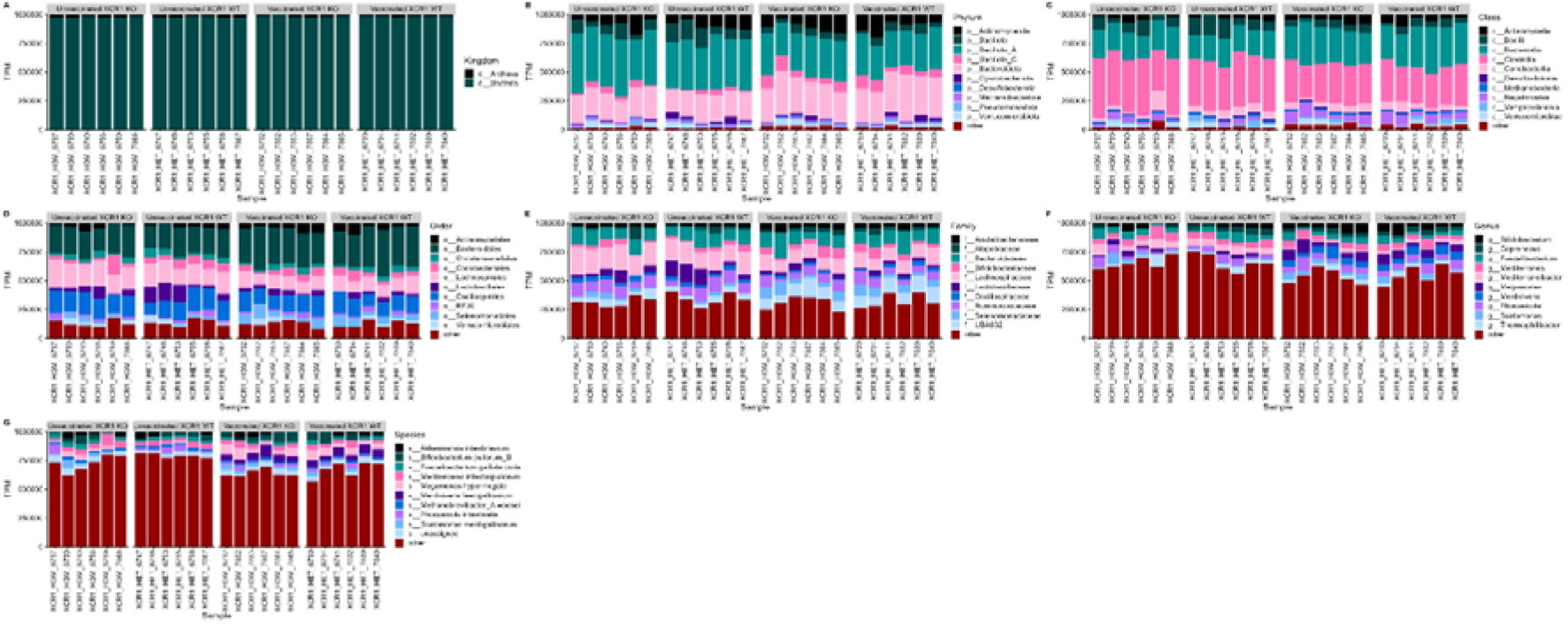
Transcripts per million (TPM) of taxa identified in the chicken caeca, as calculated by mapping Illumina reads to species-level MAGs. Showing the ten most abundant taxa per taxonomic level, with other taxa collected under “other”. XCR1 WT: heterozygous for the XCR1-iCaspase9-RFP transgene (wild-type for XCR1 expression). XCR1 KO: homozygous for XCR1-iCaspase9-RFP transgene (deficient for XCR1 expression). Vaccinated: vaccinated against IBV. Unvaccinated: not vaccinated against IBV. A) Kingdom. B) Phylum. C) Class. D) Order. E) Family. F) Genus. G) Species.

**Table 3:** Metabolic functional trait profiles for species-level MAGs, as output by METABOLIC.

**Table 4:** AMR genes identified in species-level MAGs using RGI with the Comprehensive Antibiotic Resistance Database.

While MAGs provide an effective way of analysing bacteria or archaeal members of the microbiota, it can require substantially greater sequencing depth to construct MAGs from taxa with larger genomes (e.g. eukaryotes) and from rarer members of the microbiota. Tools such as Kraken2 assign taxonomic labels to sequencing reads based on a user defined database, and can thus be used to identify taxa that would not be captured by MAGs. After classifying both our HiFi and Illumina reads, a far greater proportion of HiFi reads (**Figure 4A**) were assigned a taxonomy in comparison to Illumina reads (**Figure 4B**). For both short and long reads, the vast majority of reads that were assigned a taxonomy were assigned to bacteria. As expected, due to the fact that MAGs are usually constructed from the most abundant taxa, the phyla that were most often assigned to reads matched those that were found to be most abundant in samples when mapping MAGs to reads (**Figure 4C**): Bacillota, Bacteroidota and Actinomycetota. Very few reads were assigned as Archaea (Illumina: 0.42 ± 0.10%), viruses (Illumina: 0.034 ± 0.051%) or eukaryotes (0.90 ± 0.22%).

**Figure 4:**
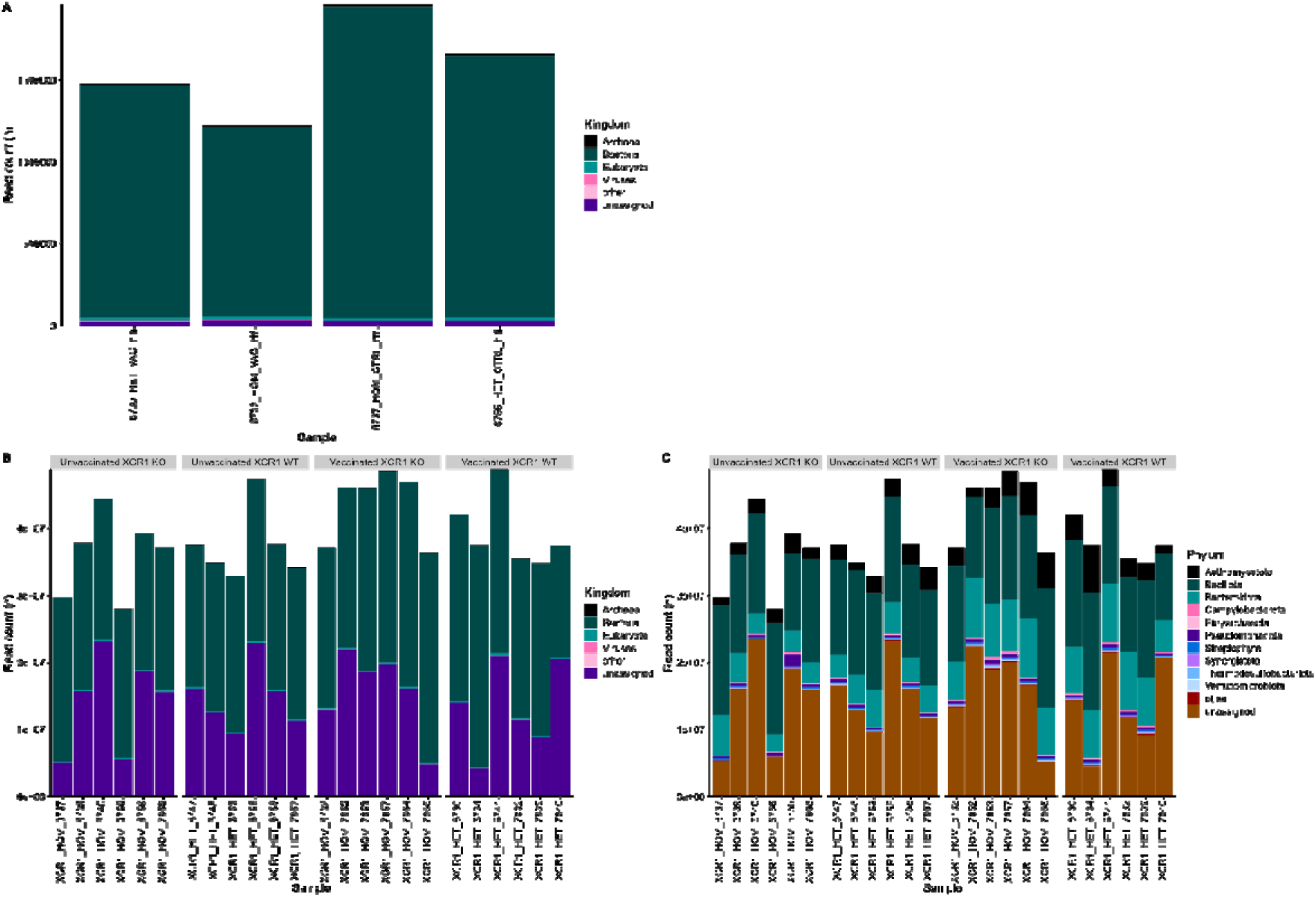
Reads taxonomically classified by Kraken2. XCR1 WT: heterozygous (HET) for the XCR1-iCaspase9-RFP transgene (wild-type for XCR1 expression). XCR1 KO: homozygous (HOM) for XCR1-iCaspase9-RFP transgene (deficient for XCR1 expression). Vaccinated/VAC: vaccinated against IBV. Unvaccinated/CTRL: not vaccinated against IBV. A) Number of PacBio HiFi reads classified by Kraken2 to each taxonomy at Kingdom level. B) Number of Illumina reads classified by Kraken2 to each taxonomy at Kingdom level. C) Number of Illumina reads classified by Kraken2 to each taxonomy at Phylum level.

### XCR1 does not affect the gut microbiota in chickens, but vaccination for IBV does

We wanted to assess whether the lack of XCR1 affected the birds’ gut microbiota compositions and whether IBV vaccination also impacted the gut microbiota. Using species-level MAG abundance data, we compared the alpha diversity (within sample diversity) between the four groups: vaccinated XCR1 WT, vaccinated XCR1 KO, unvaccinated XCR1 WT, and unvaccinated XCR1 KO. No significant differences were observed between groups for diversity (**Figure 5A**, Inverse Simpson’s: p = 0.1). However, richness was found to differ significantly between groups (**Figure 5B**, Chao1 index: p = 0.00752). After conducting pairwise comparisons of richness, only vaccinated XCR1 KO and unvaccinated XCR1 KO birds were found to differ significantly (Chao1 index: p = 0.026). That we found significant differences in richness but not diversity between our samples indicates that while there are differences in the numbers of species between groups, the evenness of the microbial communities does not differ significantly. The inverse Simpson’s index gives more weight towards common members of the microbiota whereas the Chao1 index is calculated based on rare members of the microbiota; likely indicating that the differences in richness between groups are due to rarer taxa.

**Figure 5:**
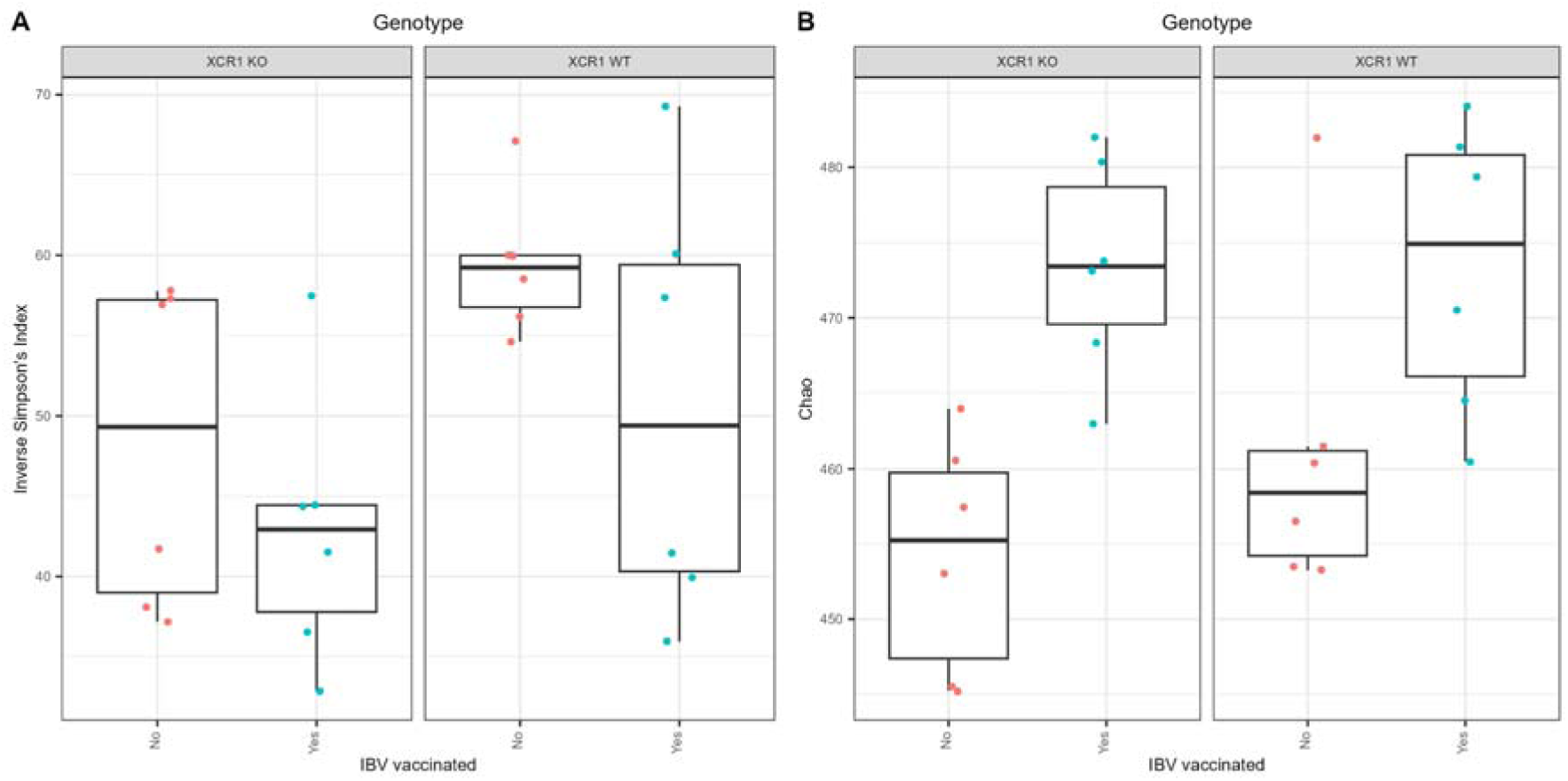
Boxplots showing diversity (A: Inverse Simpson’s index) and richness (B: Chao1 Index) of the caecal microbiota. XCR1 WT: heterozygous for the XCR1-iCaspase9-RFP transgene (wild-type for XCR1 expression). XCR1 KO: homozygous for XCR1-iCaspase9-RFP transgene (deficient for XCR1 expression).

Beta-diversity measures the similarity of microbiota compositions between samples. PERMANOVA was used to test differences in beta-diversity (Bray-Curtis) between groups, with sex, IBV vaccination, and genotype included as covariants, and including interactions between these groups. Chickens of different sexes did not have significant differences in their gut microbiota composition (p = 0.1). However, both genotype (p = 0.043) and vaccination status (p ≤ 0.00001) were found to correlate with significant differences in the gut microbiota composition (**Figure 6**). No significant interactions between covariants were identified.

**Figure 6:**
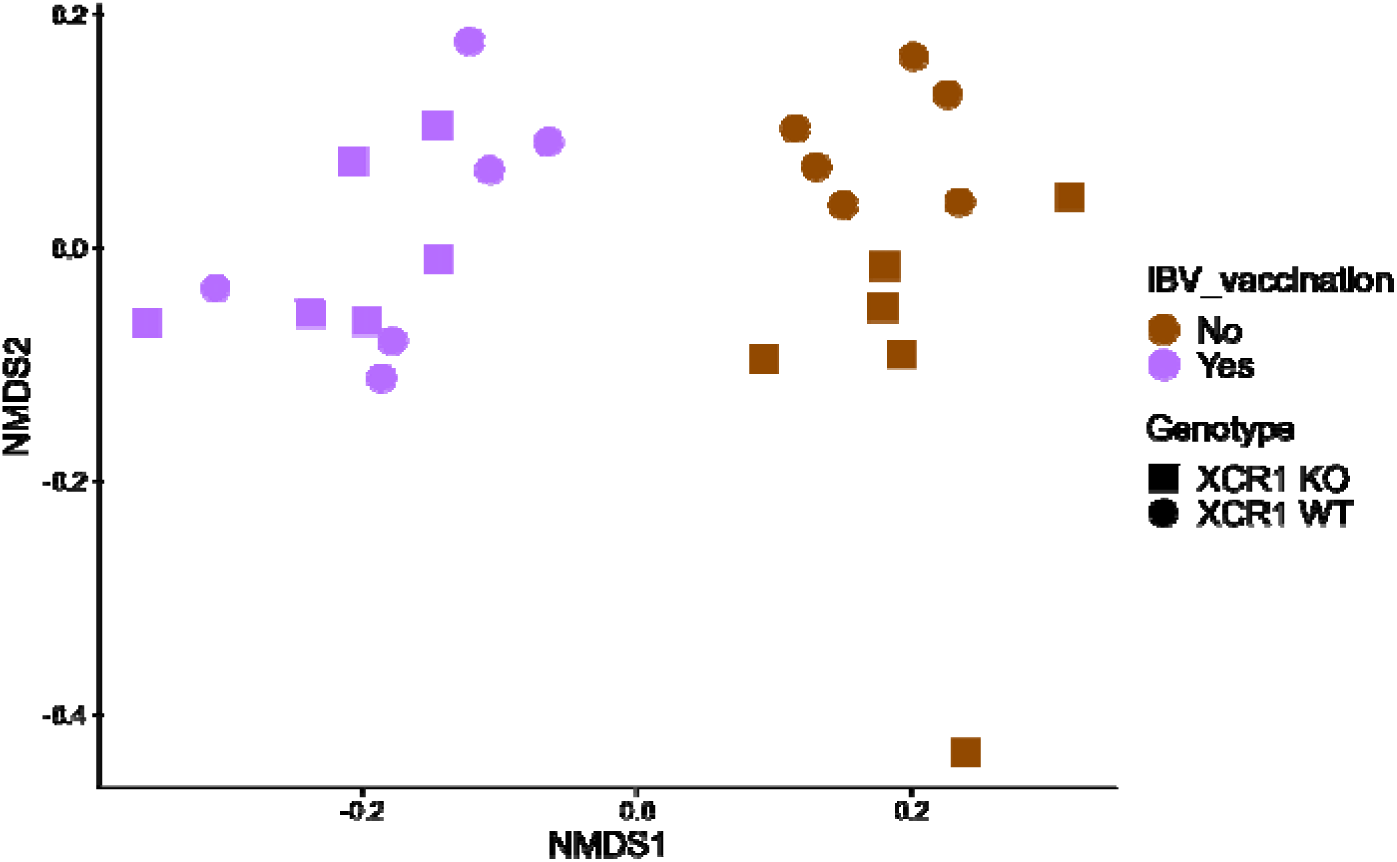
NMDS clustering samples using Bray-Curtis dissimilarity values (stress=0.11). XCR1 WT: heterozygous for the XCR1-iCaspase9-RFP transgene (wild-type for XCR1 expression). XCR1 KO: homozygous for XCR1-iCaspase9-RFP transgene (deficient for XCR1 expression).

ANCOM-BC2 was used to identify individual species-level MAGs and genera that differed between groups, with sex, genotype and vaccination status as covariants. Only one MAG was identified as being differently abundant between sexes: metabat2_coassembly_bwa.518 (p = 0.02, *Ornithomonoglobus_A intestinigallinarum).* The genus to which this MAG belongs was also identified as being significantly differently abundant between sexes (p = 0.01). However, this MAG is at a very low abundance in our samples; its maximum relative abundance in our Illumina samples is 0.07%, therefore it is likely that this difference is not biologically relevant. No species-level MAGs or genera were found to differ significantly based on genotype. However, vaccination status was found to be significantly correlated with changes in numerous taxa, including 95 species-level MAGs and 40 genera (**Figure 7**). Of the twenty-five genera that were significantly more-abundant in the IBV unvaccinated group, the majority were members of the Bacillota phylum (n=20), with the remaining genera belonging to the phyla Actinomycetota (n=3), Bacteroidota (n=1) and Cyanobacteriota (n=1). Conversely, of the fifteen genera that were significantly more-abundant in the IBV vaccinated group, only one belonged to the phylum Bacillota (*UMGS1491*). The remaining fifteen genera originated from a wide diversity of phyla, including Actinomycetota (n=4), Bacteroidota (n=3), Cyanobacteriota (n=2), Deferribacterota (n=1), Pseudomonadota (n=3), Verrucomicrobiota (n=1).

**Figure 7:**
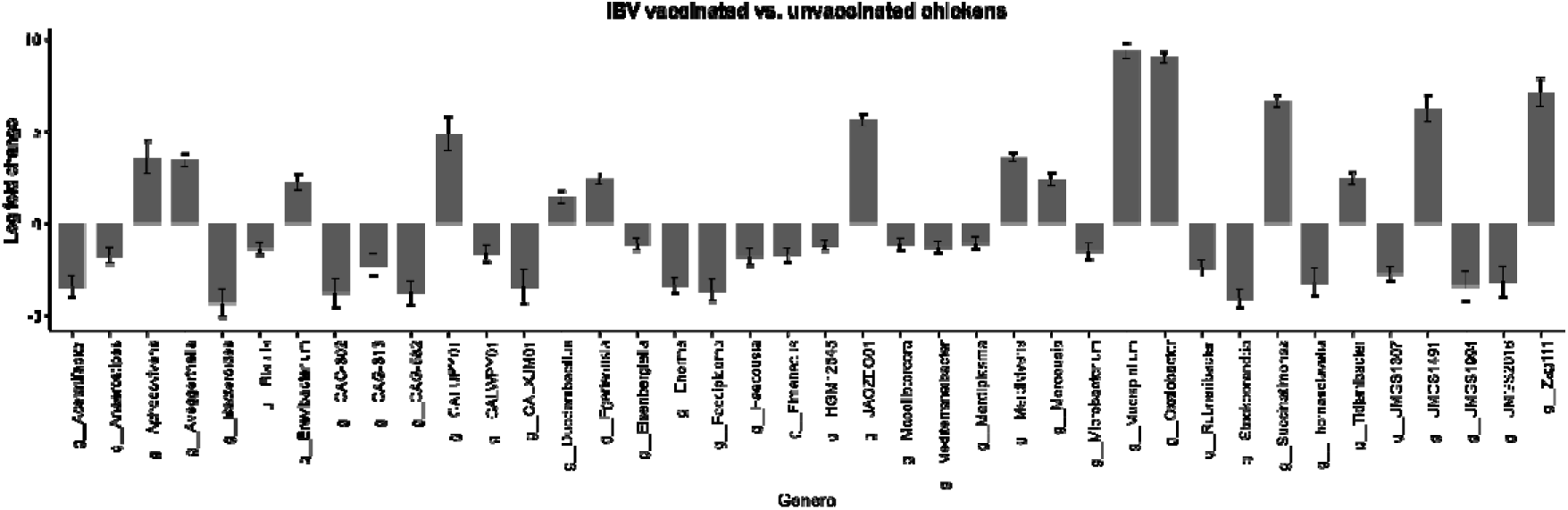
Genera that were found to be significantly different in caecal contents between birds that had or had not received an IBV vaccination. Shown as log10 fold change in TPM between IBV vaccinated vs unvaccinated chickens. Positive log fold changes indicate that the genus was more abundant in vaccinated birds, negative log fold changes indicate that the genus was more abundant in non-vaccinated birds

## Discussion

While XCR1 is known to play an important role in gut homeostasis and microbiota tolerance, its role in modulating the microbiota in chickens has not previously been characterised. Using long-read (PacBio HiFi) and short-read metagenomic sequencing, we compared the caecal microbiota of wild-type and XCR1 knockout chickens, that were or were not vaccinated for IBV.

A combined short and long read sequencing approach was taken, in order to produce higher-quality MAGs that would be of greater use as reference genomes, while also producing MAGs from less abundant members of the gut microbiota. Short-read sequencing is a cost-effective method for MAG construction, and it is therefore possible to sequence microbiota samples to a high depth and to capture rarer members of the microbiota. However, this method rarely produces complete genomes that are of a high-enough quality to be used as reference genomes (67). Alternatively, long-read sequencing can be used to produce high-quality, complete genomes from metagenomic data, including data from the chicken gut (27, 68).

In this study, we constructed 480 species-level MAGs, 74 of which were single contig MAGs produced from either long-read or hybrid assembly. As shown previously (69), we found that reads produced by HiFi sequencing were more likely to be taxonomically classified than Illumina reads, most likely due to their increased length. As with previous studies, one of the most abundant phyla in the chicken caeca, both in terms of number of MAGs and in relative abundance within samples, was Bacillota (16, 68, 70). Our samples also contained a high proportion of Bacteroidota, which is rare in young birds not exposed to an adult gut microbiota, but is more common in maternally raised birds and in older birds (71–73). This may be due to our birds being older than is standardly used in chicken microbiota studies. At sampling our birds were 19 weeks old, and it is common in chicken microbiota research to study birds that are 0-6 weeks of age, as this corresponds to the lifespan of a commercial broiler bird. Very few of our reads were taxonomically assigned to fungi, viruses, archaea and eukaryotes, which agrees with previous findings (16, 70).

When comparing wild-type and XCR1 knockout birds we found few differences in the gut microbiota composition: while beta-diversity was found to be significantly different between the genotypes this effect was subtle, with vaccination having a far greater impact on overall microbiota community composition. No differences in alpha-diversity or in the abundance of specific MAGs and genera were found. These findings are surprising, as similar mouse models have demonstrated that XCR1 plays an important role in gut homeostasis (13). It is possible that under a healthy-state there is minimal effect on the gut microbiota due to lack of XCR1, but that this impact may be increased when the gut environment is subjected to perturbations, as mice that are deficient in XCR1+ cDCs do not exhibit dysregulated gut inflammation during a steady-state but are more susceptible to chemically-induced colitis (9, 13). It is also possible that exposing chickens to different microbial sources early in life may lead to differences in the relationship between cDCs and the microbiota, as the gut microbiota plays an important role in shaping the chicken immune system (2), with early-life antibiotic exposure; exposure to adult caecal microbes and raising environment (cage vs. litter floor pens) all impacting the maturation of the chicken immune system (4, 74, 75).

It has also previously been observed that depletion of the microbiota leads to an impaired immune response to nephropathogenic IBV (76). In our study, large, genotype-independent differences in the composition of the microbiota were observed in IBV vaccinated vs. unvaccinated birds. Birds differed both in terms of the beta-diversity and richness of their microbiota, with vaccinated birds having richer caecal microbiota communities. This differs from a previous study by Borey *et al.* using 16S rRNA gene sequencing to examine IBV vaccination effect in several inbred chicken lines after the birds had also been challenged with IBV infection (19). Alongside suggesting a link between the immune response to vaccination and the microbiota (involving TCR□δ+ T cells), this study found opposite results to our own: that vaccinated birds had decreased microbiota richness in comparison to unvaccinated birds. Also, none of the operational taxonomic units identified as significantly differing between vaccination groups in this previous study originated from genera that were found to differ between vaccination groups in our study, except for *Eisenbergiella* which in the previous study was found to be significantly more abundant in vaccinated birds but in our study was found to be significantly less abundant. These differences between studies may be caused by several factors, including the fact that in our study birds were only given an IBV vaccination, whereas Borey *et al.* also challenged birds with IBV infection. It has previously been shown that IBV infection alone is associated with differences in the gut microbiota composition (77–79). Differences may also be due to rearing conditions or genotype, both of which have been shown to significantly affect the chicken gut microbiota (80, 81).

It should be noted when interpreting our results that we are not able to separate the possible effect of penning from that of IBV vaccination. While all birds were initially housed in the same room, vaccinated birds were moved to a different room immediately prior to vaccination as the vaccination used contains a live strain that could be transmitted via aerosols. Borey *et al.* also housed vaccinated vs. unvaccinated birds separately, and it is therefore also not possible to separate these effects from their study (19). Pen has previously been shown to significantly affect the composition of the chicken microbiota (80), as has stress (82), which may result from birds being moved to a new environment, though these observed effects were subtler than those observed in our study.

In conclusion, we found that in a healthy state a lack of XCR1 on chicken cDCs is not related to large changes in the microbiota of chickens. This may differ in birds challenged with gut injury or infection, and this would be an interesting avenue for future research. IBV vaccination may be related to larger changes in the chicken gut microbiota structure, but these results should be taken with caution due to the potential for housing effects.

## Ethical approval and consent to participate

All birds were hatched and housed in premises licensed under a UK Home Office Establishment License in full compliance with the Animals (Scientific Procedures) Act 1986 and the Code of Practice for Housing and Care of Animals Bred, Supplied or Used for Scientific Purposes. Production of the XCR1 knockout line (10) was carried out under UK Home Office Licenses (70/8528; 70/8940 and PP9565661). Ethical approvals were obtained from The Roslin Institute’s and University of Edinburgh’s Animal Welfare Ethics Review Board and conducted under the authorisation of a UK Home Office Project License (PE263A4FA) and adhered to the guidelines and regulations of the UK Home Office ‘Animals (Scientific Procedures) Act of 1986’.

## Author contributions

Laura Glendinning contributed to conceptualization, writing – original draft, writing – review & editing, investigation, visualization, formal analysis, and data curation. Zhiguang Wu contributed to investigation, project administration and writing – review & editing. Mick Watson contributed to conceptualization, funding acquisition, and writing – review & editing. Lonneke Vervelde contributed to investigation, and writing – review & editing. Adam Balic contributed to conceptualization, funding acquisition, investigation, project administration, supervision, and writing – review & editing.

## Acknowledgements

We would like to thank the staff at the National Avian Research Facility for the care of our animals. PacBio Sequel IIe sequencing was carried out by Edinburgh Genomics, The University of Edinburgh.

